# Global Biogeography of Prokaryotes in Mangrove Sediments: Spatial Patterns and Ecological Insights from 16S rDNA Metabarcoding

**DOI:** 10.1101/2024.12.19.629522

**Authors:** Emma Jamon-Haon, Philippe Cuny, Alix Rossi, Léa Sylvi, Maud Fiard, Cécile Militon

**Affiliations:** Aix Marseille Univ, Université de Toulon, CNRS, IRD, MIO, Marseille, France

**Keywords:** Mangroves, prokaryotes, biogeography, 16S rDNA metabarcoding, sediments

## Abstract

The global asymmetry in the floristic diversity distribution within mangrove ecosystems is well-documented. However, sediment microbial communities, which play crucial ecological roles, are often overlooked in mangrove biogeography studies. This study aimed to investigate the diversity, composition, and distribution of prokaryotic communities across distinct mangrove bioregions (12 countries and territories from the Caribbean bioregion, Southeast America, East Africa, Indo-Malaysia and Australasia). We conducted a meta-analysis by compiling 16S rDNA datasets from 17 previous studies (540 samples) over a six year period (2014-2020) and applied community ecology analyses combined with random forest models. Although no significant correlation was observed between tree richness and microbial diversity, a microbial hotspot was identified in the Indo-West Pacific region. Microbiota associated with different mangroves displayed opposite latitudinal diversity gradients and beta-dispersions. Distance-decay relationships were weak but statistically significant. Random forest models identified key microbial taxa, and the OTUs shared across all bioregions suggested the presence of a potential core mangrove microbiota. Taxonomic and statistical analyses underscored the great heterogeneity of microbiota composition, heavily influenced by the study (e.g., molecular and bioinformatic methodologies). Future research on mangrove microbiome would benefit from standardized sampling and sequencing methods. Despite current limitations, distance-decay relationship confirmed the influence of dispersal barriers for mangrove sediment microbiota. This study also suggests that environmental factors, rather than plant diversity alone, may play a prominent role in shaping the diversity, composition and distribution of prokaryotic communities in mangrove sediments.

## Introduction

Mangroves are intertidal forests found along relatively sheltered coastal margins of intertropical regions, covering approximately 75% of these areas (Giri et al., 2011). The tree and shrub species primarily composing these forests are considered as ecosystem engineers as they trap sediments and support diverse life forms. Fossil records and molecular markers indicate that Indo-West-Pacific (IWP) and Atlantic-East-Pacific (AEP) mangroves diverged around 10 to 15 Mya, coinciding with the closure of the Tethys Seaway and the mid-Miocene cooling (Takayama et al., 2021; Harzhauser et al., 2007). Consequently, the AEP and IWP are separated by the African continent and the Pacific Ocean, with no shared tree species between them (Tomlinson, 2016). Further disjunctions within these regions have formed subregions, including Eastern Pacific, Western Atlantic, Caribbean, West Africa, East Africa, Indo-Malaysia, and Australasia (Hogarth, 1999). Excluding hybrids, the IWP region hosts 54 true mangrove species, with a biodiversity hotspot in the Indo-Malaysian bioregion, whereas the AEP region is home to only 17 species, with a hotspot in the Caribbean (Duke, 1992; Duke, 2017). This creates a significant asymmetry in the global distribution of floristic diversity in mangrove assemblages. Previous studies have shown that factors such as environmental conditions, biotic interactions, ocean currents and geographic barriers play a crucial role in the distribution of mangrove species (Duke et al, 1998). However, beyond the diversity of mangrove flora, these ecosystems also harbor rich sediment microbial communities that play essential roles in mangrove functioning, including organic matter decomposition and nutrient cycling (Lai et al., 2022). Despite their critical roles, sediment microbial communities are often overlooked in discussions of mangrove biogeography.

Compared to other biomes, mangrove sediment prokaryotic communities are particularly rich and complex, shaped by extreme environmental gradients like salinity, oxygen, and temperature, as well as high rates of bioturbation (Zhang et al., 2019; Booth et al., 2023). Additionally, climate and precipitation patterns have been shown to play key roles in shaping microbial communities (Zhang et al., 2019). Dominated by groups like Gammaproteobacteria, Deltaproteobacteria, Alphaproteobacteria, Chloroflexi, Firmicutes, Euryarchaeota and Crenarchaeota, these communities are highly sensitive to environmental heterogeneity (Lai et al., 2022). Studies from different bioregions have emphasized that tree species significantly affect mangrove sediment microbiota composition through selective processes (Gomes et al., 2013; Wu et al., 2016; Muwawa et al., 2020; Mai et al., 2021; Sui et al., 2023), with the combined actions of root exudates, litter deposition, and associated endofauna (Twilley et al., 2019; Zhuang et al., 2020). These relationships raise the question whether global patterns in microbial community distribution parallel the known global biogeographic patterns of mangrove trees.

The global distribution of microbes is shaped by mechanisms such as dispersal, environmental filtering, adaptation, diversification, and local extinction (Xu et al., 2020). The longstanding hypothesis that “everything is everywhere, but the environment selects” (Baas Becking, 1934) posits that microbial distribution is determined primarily by environmental selection due to their small size and high densities, as well as their ability to enter dormancy and disperse. However, for host-associated microbes, particularly those of sessile plants, geographic structuring along environmental gradients is likely, potentially resulting in patterns like distance-decay relationships or latitudinal diversity gradients (Härer and Rennison, 2023). To date, these hypotheses regarding microbial biogeography have rarely been tested in mangrove sediments, despite these ecosystems being ideal candidates for such studies.

Recent studies have barely begun to explore microbial biogeography in mangroves. For instance, Du et al. (2023) observed country-scaled distance-decay relationships in prokaryotic communities between China and South America. Tavares et al. (2021) further demonstrated that local environmental factors in Brazil better explained microbial community differences than geographic distance, even amidst genetic divergences among mangrove trees. Similarly, Thomson et al. (2022a) found that microbial communities in Australian and Saudi Arabian mangroves, differed primarily in structure rather than composition. Building on these studies, we hypothesized that mangrove sediment prokaryotic communities are shaped by deterministic processes and may reflect the global biogeographic patterns of their host plants.

In this study, we aimed to investigate the diversity, composition, and distribution of prokaryotic communities across different mangrove bioregions on a global scale, a novel approach that has not been previously tested for these ecosystems. To achieve this, we leveraged the extensive pool of 16S rDNA metabarcoding data generated in recent decades to investigate the composition of the prokaryotic community in mangrove sediments on a global scale (Lai et al., 2022). Metabarcoding is one of the most widely used methods for assessing the environmental biodiversity of microbes, organisms invisible to the naked eye and still mostly uncultured.This interest has been accompagnied by the diversification of molecular techniques to study their biodiversity (e.g. extraction methods, primer choice) and bioinformatical pipelines for sequence processing (Compson et al., 2020). Recognizing the variation in molecular and bioinformatic methodologies across the studies, we developed a Python algorithm to standardize the analysis by targeting the V4 region of the 16S rDNA gene (540 samples from 17 studies). Using community ecology analyses and random forest models (Cutler et al., 2007), we identified key microbial taxa and their potential ecological drivers, contributing to our understanding of microbial biogeography in mangrove ecosystems.

## Methods

### 16S rDNA datasets collection

A search was conducted on the NCBI Sequence Reads Archive (SRA) on April, 18 2023, using the query « mangrove* AND sediment* » (see PRISMA diagram in Supplementary Figure 1, Haddaway et al 2022). A total of 5,285 SRA runs (*i.e.* files related to one sequenced sample in SRA format) from 110 bioprojects (*i.e.* collections of biological data related to a research effort) were extracted, meeting two main criteria: i) they were in fastq format and ii) the samples were sequenced using Illumina platforms. After extracting the runs from SRA, they were filtered based on SRA metadata and supplemented with additional searches in related publications (when available). Briefly, we excluded runs from samples that were not assigned to the correct bioproject, did not target the 16S rRNA gene, were not from sediments, lacked a related published paper, did not have available geographical coordinates , or involved anthropogenically altered mangroves, cultivation approaches, or forced bioturbation. We retained only the runs targeting a portion of the 16S rRNA gene containing the V4 region and collected between 0 cm and 20 cm, as these dominated the dataset. At this stage, 904 runs from 33 related studies with a corresponding DOI remained. These runs were not deposited in a consistent format across the different bioprojects (*e.g.* missing primers, absent R1 or R2 files, pre-merged sequences, or no corresponding sample names). A standardization effort was made to recover raw sequencing files or obtain further information about the runs by contacting the authors. At this process, runs from 24 studies were ready for preprocessing. Metadata were compiled based on SRA metadata, literature searches, or directly correspondence with authors, and included only qualitative variables such as bioregion, mangrove geomorphological type, country, related study, or primers used.

### Processing of the reads and OTU clustering

To obtain comparable data, we developed a Python script using a regular expression to identify all sequences matching a consensus region with a one-base mismatch tolerance, corresponding to the universal primers 515F (in the R1 fastq files) and 806R (in the R2 fastq files) (5’-GTGYCAGCMGCCGCGGTA-3’ and 5’-GACTACNVGGGTWTCTAAT-3’, respectively). The script then generates a new dataset in fastq format, containing only the bases downstream of the *in silico* primers. If the primer patterns were not found in a sequence, that sequence was excluded from the new file, resulting in orphaned sequences lacking complementary sequences in the other file, which were also discarded. The reads were then processed per study following the standard pipeline of ‘dada2’ 1.24.0 (Callahan et al., 2016). The ‘truncLen’ argument was adjusted based on the length of the reads (250 or 300 b) and the observed diminishing quality scores in each study. Amplicon Sequence Variants (ASVs) were taxonomically assigned with SILVA 138.1 database (Quast *et al*., 2012), and the sequences corresponding to chloroplasts and mitochondria were removed. We then clustered ASVs into Operational Taxonomic Units (OTUs) using a 99% similarity threshold (Edgar, 2018) to minimize variation arising from the different runs of sequencing with the function ‘asv2otu’ from ‘MiscMetabar’ 0.40. At this stage, some samples were removed due to data corruption (*e.g.* could not be read by the ‘filterAndTrim’ function for unknown reason) or insufficient sequencing depth. Additionnally, studies with less than five samples were removed. Ultimately, we retained 17 studies, resulting in 540 samples.

### Taxonomy and statistical analyses

Most abundant taxa were displayed using the ‘fantaxtic’ package (Teunisse, 2022). Alpha-diversity was computed on rarefied samples to account for uneven sequencing depths across samples and studies, which involved randomly drawing 3,000 sequences from each sample. The computed taxonomy indices included OTU richness (S) and Shannon-Weaver diversity (H’). Significant differences were tested between qualitative variables. Normality was assessed using Shapiro’s tests, and homoscedasticity was evaluated with Levene’s tests. Since both conditions were not met and the number of samples per tested group was quite heterogeneous, significant differences between categories that involved two groups (*e.g.* AEP versus IWP) were tested non-parametrically using Mann-Whitney test, while Dunn’s pairwise multiple comparisons were conducted for parameters involving more than two groups (*e.g.* bioregion, mangrove type, study). The skewness of the distributions was assessed using the ‘datawizard’ package (Patil *et al*. 2022). We calculated Spearman’s correlations to evaluate the effects of latitude and longitude on alpha-diversity using the cor.test function from the ‘stats’ package, which implements algorithm AS 89. The distance-decay relationship was tested with a linear regression, with the data from Thomson et al (2022a) excluded due to the compositional similarity between the Saudi Arabian and Australian samples, which was influenced by methodological factors (see the results).

Beta-diversity was visualized by computing non-metric multidimensional scaling (NMDS) on the Jaccard distance matrix with ‘metaMDS’ from ‘vegan’ v2.6-4. The effects of qualitative variables on beta-diversity were tested using Permutational Analysis of Variance (PERMANOVA) using ‘adonis2’ (1000 permutations) on the Jaccard distance matrix. Given that community dissimilarities were significantly influenced by the methodological parameters and varied according to the original study, we employed a random forest (RF) model approach to uncover the importance of bioregions (variables) and taxa (features) (Cutler *et al*., 2007). To reduce the high dimensionality of the OTU table and mitigate the dominance of zeros, we selected only the most abundant OTUs (> 1% in at least one sample). Proportional abundances of OTUs were retained for classification (ranging between 0 and 1). RF models were computed using ‘scikit-learn’ v1.5.0 in Python (Pedregosa *et al*., 2011). A classifier model was constructed for each parameter based on a training set (80%) and tested on a separate set (20%). To avoid overfitting, each model was built with 15 estimators, a maximum depth of 8, and the Gini coefficient as decision criterion. Model scores were computed using a 10-fold cross-validation. OTU importances, based on the mean accumulation of impurity decrease within each tree, were retrieved and ranked for each model of the cross-validation. OTUs with non-zero importance in at least 8 out of 10 models were considered the most important OTUs for the corresponding class.

## Results

A total of 540 samples from 12 countries and territories were analyzed in the study: 130 from the Caribbean bioregion, 40 from Southeast America, 169 from East Africa, 154 from Indo-Malaysia and 48 from Australasia (Figure 1). The number of mangrove tree species reported in the studies varied between bioregions, with 3 species from 3 genera in AEP and 12 species from 7 genera in IWP. The sampling periods spanned from 2014 to 2020, with most samples collected outside of the rainy seasons (Table 1). Coastal and estuarine mangroves constituted over 72% of the samples.

**Figure 1.**
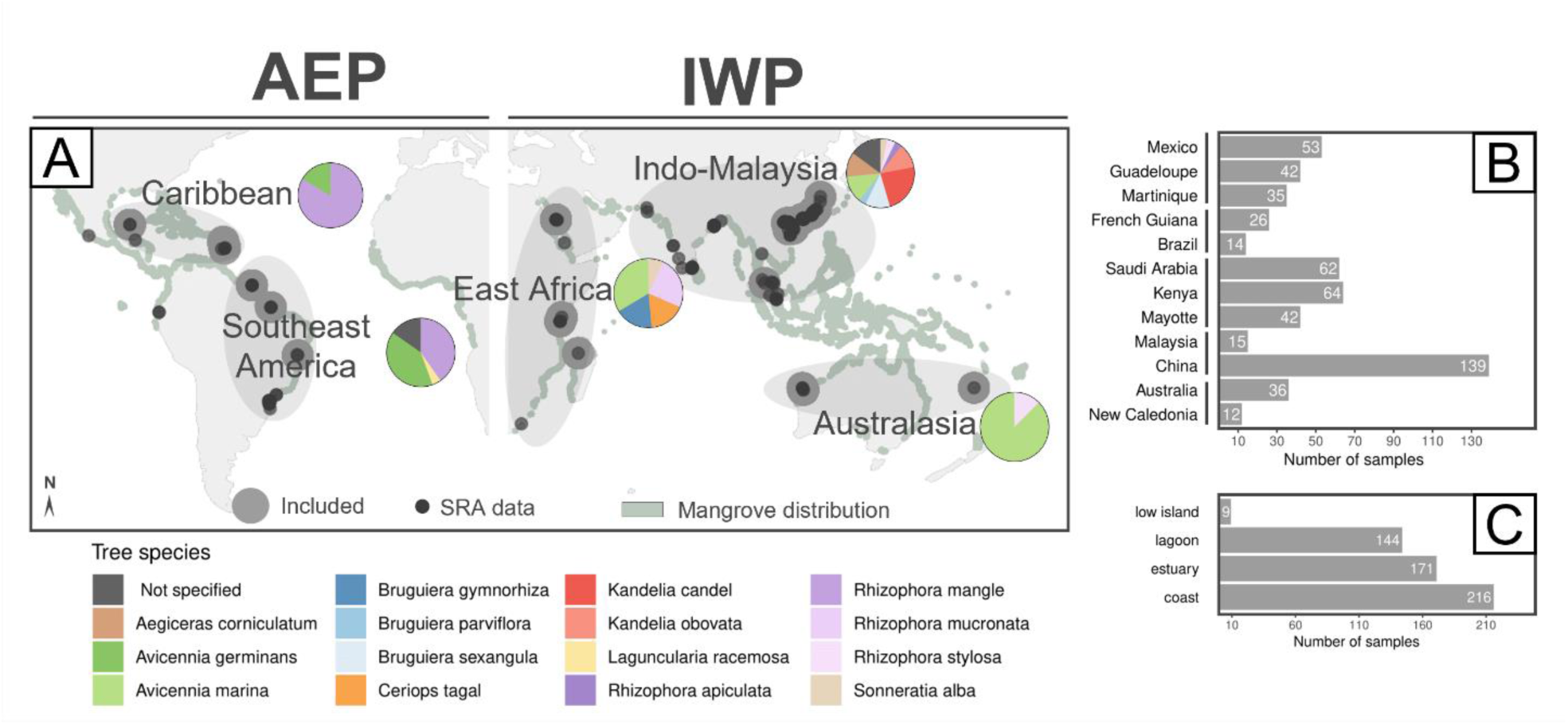
**(A)** Map of sampling locations of data retrieved on SRA that covered 16S regions from mangrove sediments (black dots), of the samples included for the meta-analysis (grey dots) and relative proportions of sample related mangrove tree species across the bioregions. Mangrove cover data (greyish green) was uploaded from Giri et al. (2011). Bar plots indicate **(B)** the number of samples per country and **(C)** for each mangrove geomorphological type.

**Table 1.** Studies included in the meta-analysis and their related production information. Letters indicate institutes of provenance. See supplementary table for more details about the metadata and the primer sequences.

| Region | SRA bioproject | Number of samples | Sampling period | Sampling depths | Extraction kit | Targetted region | Forward primer | Reverse primer | Taq polymerase |
| --- | --- | --- | --- | --- | --- | --- | --- | --- | --- |
| **Atlantic-East-Pacific (AEP)** |  |  |  |  |  |  |  |  |  |
| **Caribbean** |  |  |  |  |  |  |  |  |  |
| Mexico | PRJNA550111<sup>a</sup> (Gómez-Acata, Teutli et al 2023) | 53 | October 2018 | 0-2 cm | DNeasy PowerSoil | V4V5 | 515F Caporaso | 806R Caporaso | Takara ExTaq |
| Guadeloupe | PRJNA1118708<sup>b</sup> | 42 | June 2019 | 0-2 cm, 2-10 cm, 10-20 cm | DNeasy PowerSoil | V4V5 | 515FY Parada | 909R WangQian | Pfu Promega |
| Martinique | PRJNA874590<sup>b</sup> (Fiard et al 2024) | 35 | June 2018 | 0-2 cm, 2-10 cm, 10-20 cm | DNeasy PowerSoil | V4V5 | 515FY Parada | 909R WangQian | Pfu Promega |
| **Southeast America** |  |  |  |  |  |  |  |  |  |
| French Guiana | PRJNA735070<sup>b</sup> (Fiard et al 2022) | 26 | November 2017 | 0-2 cm, 2-10 cm, 10-20 cm | DNeasy PowerSoil | V4V5 | 515FY Parada | 909R WangQian | Pfu Promega |
| Brazil | PRJNA608697<sup>c</sup> (De Santana et al 2021) | 9 | July 2018 | 0-10 cm | MO BIO PowerSoil | V4V5 | 515FY Parada | 806R Caporaso | KAPA Hifi |
| Brazil | PRJNA817610<sup>d</sup> (Costa et al 2023) | 5 | January 2018 | 0-20 cm | MO BIO PowerSoil | V3V4 | Bakt_341F Herlemann | Bakt_805R Herlemann | KAPA Hifi |
| **Indo-West Pacific (IWP)** |  |  |  |  |  |  |  |  |  |
| **East Africa** |  |  |  |  |  |  |  |  |  |
| Mayotte | PRJNA1118920<sup>b</sup> | 42 | October 2018 | 0-2 cm, 2-10 cm, 10-20 cm | DNeasy PowerSoil | V4V5 | 515FY Parada | 909R WangQian | Pfu Promega |
| Kenya | PRJNA644929<sup>e</sup> (Muwawa et al 2021) | 20 | May 2018 | 1-5 cm, 10-15 cm | DNeasy PowerSoil | V3V4 | Bakt_341F Herlemann | Bakt_805R Herlemann | High Fidelity DNA Q5 |
| Saudi Arabia | PRJNA720541<sup>f</sup> (Thomson et al 2022a) | 62 | October 2018 | 0-2 cm, 5-10 cm | DNeasy PowerSoil | V3V4 | Pro341F Takahashi | Pro805R Takahashi | - |
| **Indo-Malaysia** |  |  |  |  |  |  |  |  |  |
| China | PRJNA692026<sup>g</sup> (Huang et al 2021) | 5 | June 2021 | 0-5 cm | Spin FastDNA | V3V4 | 338F Huws | 806R Caporaso | FastPfu |
| China | PRJNA732523<sup>h</sup> (Liu et al 2022) | 54 | April 2020 | 5-15 cm | HiPure sediment DNA Kit B | V3V4 | Bakt_341F Herlemann | Bakt_805R Herlemann | - |
| China | PRJNA475455<sup>i</sup> (Zhang et al 2019) | 36 | September 2017 | 0-10 cm, 10-20 cm | DNeasy PowerSoil | V4 | 515F Caporaso | 806R Caporaso | Takara ExTaq |
| China | PRJNA797991<sup>j</sup> (Zhang et al 2022) | 6 | November 2020 | 0-20 cm | Spin FastDNA | V4 | 515FY Parada | 806R Apprill | Phusion Hifi |
| China | PRJNA404001<sup>k</sup> (Zhu et al 2018) | 20 | June 2014 | 0-5 cm | Spin FastDNA | V4 | 515F Caporaso | 806R Caporaso | Phusion Hifi |
| China | PRJNA779243<sup>l</sup> (Zhu et al 2022) | 18 | ? 2020 | 0-15 cm | DNA extraction TiangenBiotechCo | V3V4 | 338F Huws | 806R Caporaso | Takara Taq |
| Malaysia | PRJNA756333<sup>m</sup> (Mai et al 2021) | 15 | December 2019 | 0-5 cm | Soil DNA Kit EZNA | V3V4 | 338F Huws | 806R Caporaso | FastPfu |
| **Australasia** |  |  |  |  |  |  |  |  |  |
| New Caledonia | PRJEB25548<sup>n</sup> (Luis et al 2019) | 12 | March and November 2016 | 0-10 cm | MO BIO PowerSoil | V4V5 | 515FY Parada | 909R WangQian | Taq Invitrogen |
| Australia | PRJNA720541<sup>f</sup> (Thomson et al 2022a) | 36 | June 2018 | 0-2 cm, 5-10 cm | DNeasy PowerSoil | V3V4 | Pro341F Takahashi | Pro805R Takahashi | - |

**a** Instituto de Ecología UNAM; **b** Mediterranean Institute of Oceanography (MIO); **c** New York University; **d** Universidade Federal do Pará; **e** Pwani University; **f** King Abdullah University of Science and Technology (KAUST); **g** College of Oceanology and Food Science, Quanzhou Normal University; **h** Sun-Yat-Sen University; **i** Shenzhen University; **j** Xiamen University Tan Kah Kee College; **k** Yantai Institute of Coastal Zone Research, Chinese Academy of Sciences; **l** Guangdong Ocean University; **m** South China Sea of Oceanography; **n** UMR CNRS 5557 Ecologie Microbienne

### Diversity patterns

A total of 128,196 total OTUs were initially identified, of which 84,293 remained after rarefaction. Observed OTU richness (S) differed significantly across both global areas and bioregions (Figure 2A). The IWP bioregion generally exhibited higher richness compared to the AEP region (mean S: 1,372 OTUs versus 1,289 OTUs). The Indo-Malaysian bioregion showed the highest richness and variability (max S: 2,024 OTUs), while Southeast America and Australasia exhibited the lowest richness (min S: 217 OTUs). Shannon’s diversity (H’), with an overall mean of 6.7±0.5 (from 3 to 7.4), followed a similar pattern to richness S (Figure 2A). However, East African samples did not differ significantly in diversity from those in Indo-Malaysian samples (mean H’: 6.8 ± 0.3 and 6.7 ± 0.5 respectively). Both AEP and IWP exhibited negative skewness in richness, indicating left-skewed distributions, indicating that lower richness was less common (Skewness: -1.2 ± 0.2 and -0.24 ± 0.2, respectively). The skewness of IWP’s richness was closer to 0, suggesting a more uniformly distributed richness. Shannon’s diversity distributions were similarly skewed, particularly for AEP (AEP Skewness: -2.7 ± 0.2; IWP Skewness: -1.4 ± 0.2). Overall, alpha-diversity distributions suggested that AEP samples were poorer and less evenly distributed than those from IWP, with less heterogeneity but more outliers.

**Figure 2.**
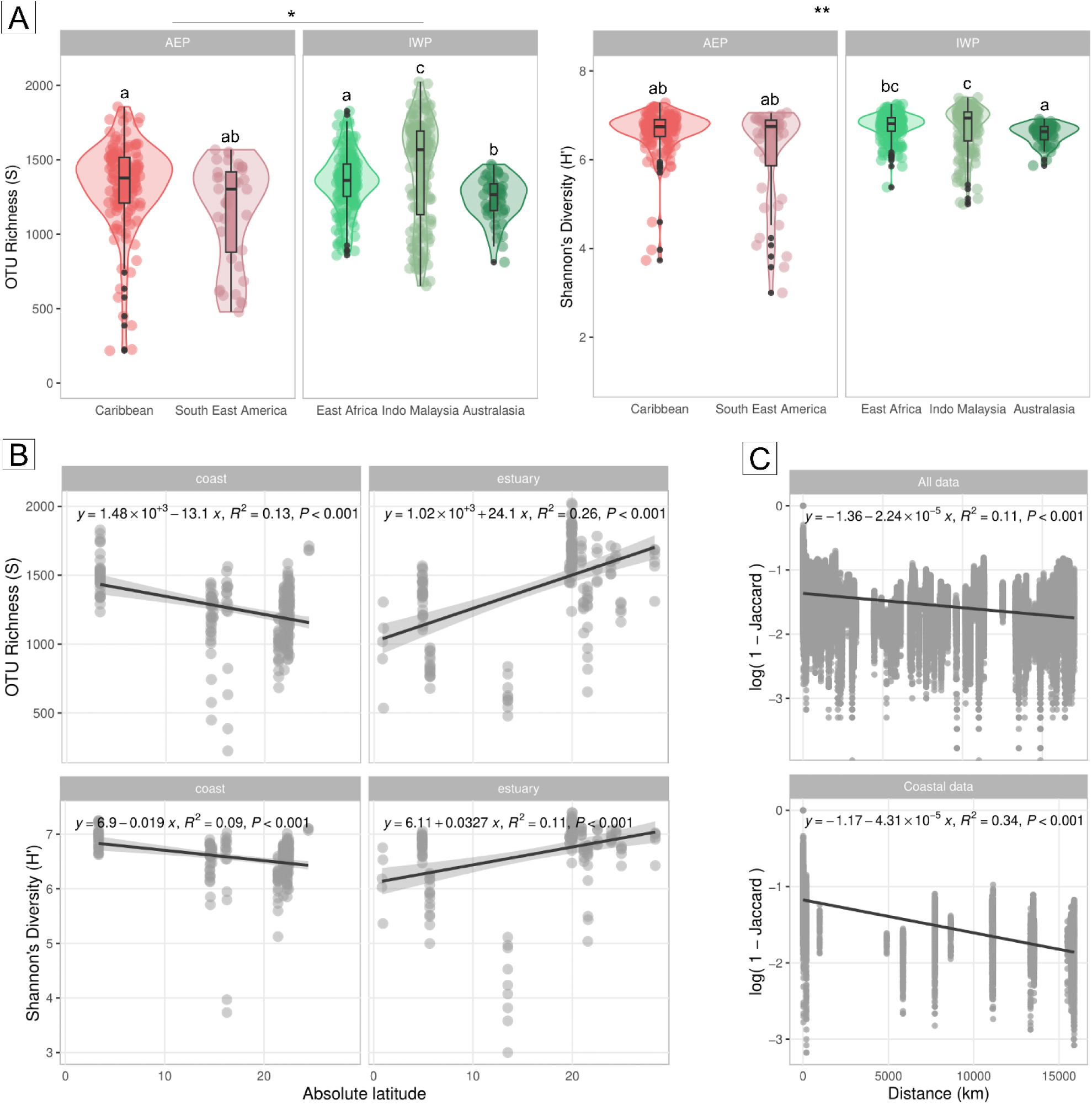
**(A)** Violin diagrams of OTU richness (left) and Shannon’s Diversity (right) in each bioregion. Letters indicate the significances of the Dunn’s pairwise comparisons (W: * p < 0.05; ** p < 0.01). **(B)** Linear regressions of OTU Richness (top) and Shannon’s Diversity (bottom) against the absolute latitude (distance from the Equator), analyzed by mangrove geomorphological types. **(C)** Distance-decay relationships without Thomson et al 2022 data, shown for all samples (top) and coastal data only (bottom).

There was no significant linear relationship between geographical coordinates and overall alpha-diversity. However, absolute latitude showed a weak but significant negative correlation with Shannon’s diversity (Spearman; H’ p < 0.0001, corr = -0.15; Figure 2B), while longitude was positively correlated with richness and diversity (Spearman; S p < 0.01, corr = 0.12; H’ p < 0.0001, corr = 0.14). Similarly, no clear relationship was found between microbial richness (S, H’) and tree richness (Supplementary Figure 2).

Focusing specifically on coastal mangroves, a significant linear relationship was observed between richness and latitude (LM; R² = 0.13, p < 0.001 for S, and LM; R² = 0.09, p < 0.001 for H’), indicating lower OTU richness and diversity further from the equator (Figure 2B). Estuarine mangroves showed the opposite trend, with higher richness and diversity away from the equator (LM; R² = 0.26, p < 0.001 for S, and LM; R² = 0.11, p < 0.001 for H’). The distance-decay relationship across all mangrove samples was weak (LM; R² = 0.12, p < 0.001 Figure 2C), though this relationship was stronger when considering only coastal mangroves (LM; R² = 0.34, p < 0.001). Estuarine mangroves could not be evaluated due to uneven geographic distribution.

### Taxonomic composition and core microbiota

Among the 97 bacterial phyla identified, Proteobacteria was the most abundant, representing 24% of total reads analyzed, dominated by *Gammaproteobacteria* (15.6%) and *Alphaproteobacteria* (7.9%) (Figure 3A, Supplementary Table 4). Chloroflexi (11%) followed, with *Anaerolineae* (9%) and *Dehalococcoidia* (1.8%) as the major classes. Other notable phyla included Desulfobacterota (7%), with *Desulfobacteria* (3.9%) and *Desulfobulbia* (2.1%) as the dominant classes, Bacteroidota (6%), Planctomycetota (4%), Acidobacteriota (3%), Actinobacteriota (2%), Firmicutes (2%), Verrucomicrobiota (1%), Cyanobacteria (1%), Myxococcota (1%), and Calditrichota (1%). Among the 16 archaeal phyla identified, Crenarchaeota was the dominant phylum (2%), represented by *Bathyarchaeia* (1.4%) and *Nitrososphaeria* (0.3%), while Thermoplasmatota contributed 1% of the total reads, mainly from the class *Thermoplasmata* (0.7%).

**Figure 3.**
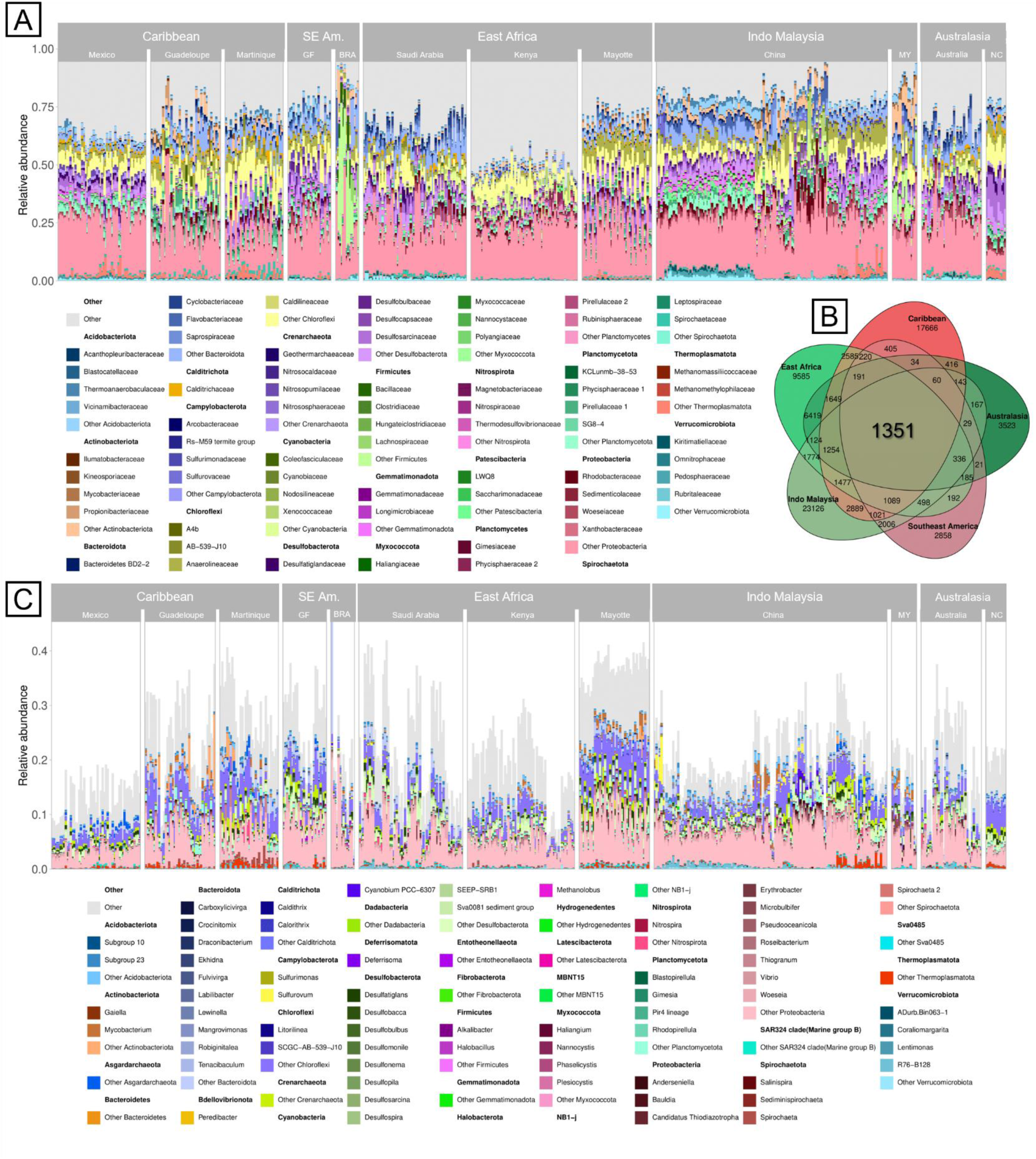
**A)** Taxonomy of top 20 Phyla and top 4 related Families observed in pooled samples from the meta-analysis of mangrove sediment microbiota. **B)** Venn Diagram illustrating the distribution of OTUs number across mangrove bioregions, highlighting 1,744 core OTUs common to all five bioregions. **C)** Taxonomy of core OTUs, showing the top genera identified for each Phylum, with relative abundances in the original samples.

A total of 1,351 OTUs (1.6%), associated with 32 phyla (Figure 3C), were shared across all five bioregions (Figure 3B), representing the OTU-based core microbiota in mangrove sediments. This core microbiota primarily comprised Proteobacteria, Chloroflexi, and Desulfobacterota (Figure 3C, Supplementary table 2). Within Proteobacteria, notable genera included *Woeseia* (10.6% of total abundance), *Thiogranum* (3.8%), *Microbulbifer* (1.4%), *Thiodiazotropha* (0.12%) and *Vibrio* (0.12%). For Chloroflexi, *Litorilinea* (3.5%) and *SCGC-AB-539-J10* (3.19%) were dominant. Desulfobacterota was represented by *Desulfatiglans* (2.4%), *Sva0081 sediment group* (2.2%), *Desulfobulbus* (0.91%), and *Desulfobacca* (0.49%). Additional genera included *Mangrovimonas* (1.1%), *Labilibacter* (0.78%) and *Lewinella* (0.72%) for Bacteroidota, and *Thermoanaerobaculaceae Subgroup 23* (2.6%) for Acidobacteriota. Other significant genera were *Mycobacterium* (1.5%) from Actinobacteria and the *Pir4 lineage* (2.6%) from Planctomycetota. Among archaea, only a few core OTUs were identified, mostly within Crenarcheaota, though none could be assigned at the genus level.

Surprisingly, a large proportion of the OTUs remained unassigned at broad taxonomic levels: 0.9% at the kingdom, 23% at the phylum, 30 % at the class, 49% at the order, 64% at the family, and up to 83% at the genus, with nearly 100% of species remaining unassigned. The proportions of unassigned OTUs were similar between bacterial and archaeal kingdoms and consistent across all the studies from which the data in this study were derived (Supplementary Table 3).

### Bioregionalmicrobiota specificities

While the main taxa were consistently shared at the phylum level across the globe, OTUs specificity varied significantly among bioregions, with only 1.6% of OTUs shared across all bioregions (Figure 3C). Between the AEP and IWP, only 21% of OTUs were shared, with AEP-specific OTUs accounting for 25% of the total and IWP-specific OTUs constituting 54%. The Indo-Malaysia bioregion contained the highest number of unshared OTUs (27%), followed by the Caribbean (21%) while Southeast America had the fewest unique OTUs (3.4%).

Non-metric multidimensional scaling (NMDS) analysis accompanied by PERMANOVA tests (Figure 4, Supplementary table 4) indicated weak but significant effects of bioregion (R²: 8.4%, Pseudo-F: 12.2, p < 0.001) and mangrove type (R²: 6.8%, Pseudo-F: 13.1, p < 0.001) on microbial community structure (Figures 4A and 4B). However, it is important to note that due to the high dimensionality and the large number of zeros in the OTU table (96.2%), the NMDS stress was quite high (> 0.2). Patchiness in microbial composition was observed at the study level, confirmed by PERMANOVA (R²: 26%, Pseudo-F: 11.7, p < 0.001) (Figure 4C).

**Figure 4.**
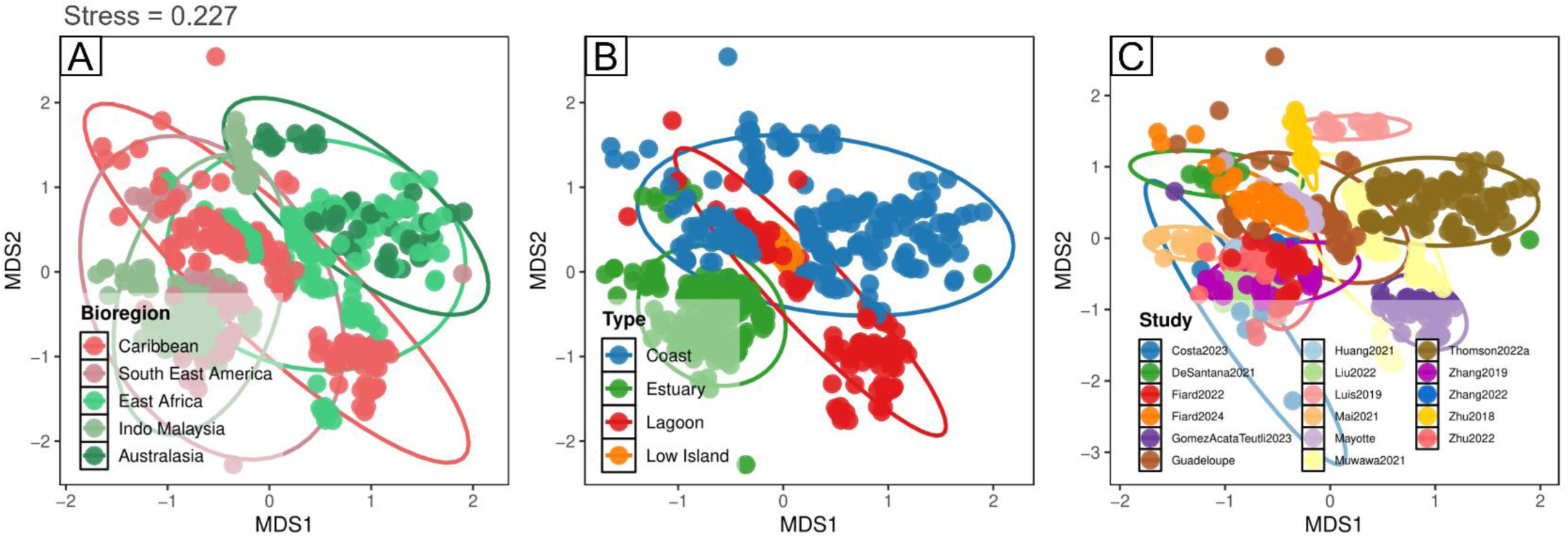
Non-metric multidimensional scaling (NMDS) plots based on Jaccard distances, colored by **(A)** bioregion, **(B)** mangrove geomorphological type, and **(C)** study.

RF models, based on 664 OTUs (filtered to include those >1% in at least one sample), achieved a bioregion classification accuracy of 90.3%. A total of 150 computed trees are presented in Supplementary Figures 4. Among the 76 most important OTUs retrieved for this model, 30 belonged to the core OTUs (Figure 5). Notably, 35 were shared with the classification model for the study class, which achieved a classification accuracy of 98%. For the mangrove type classification model, 66 OTUs were identified as being most important, resulting in a classification accuracy of 96%. Of these, 24 OTUs were shared with the bioregion model, and 33 OTUs were common with the study class model (Supplementary Table 5).

**Figure 5.**
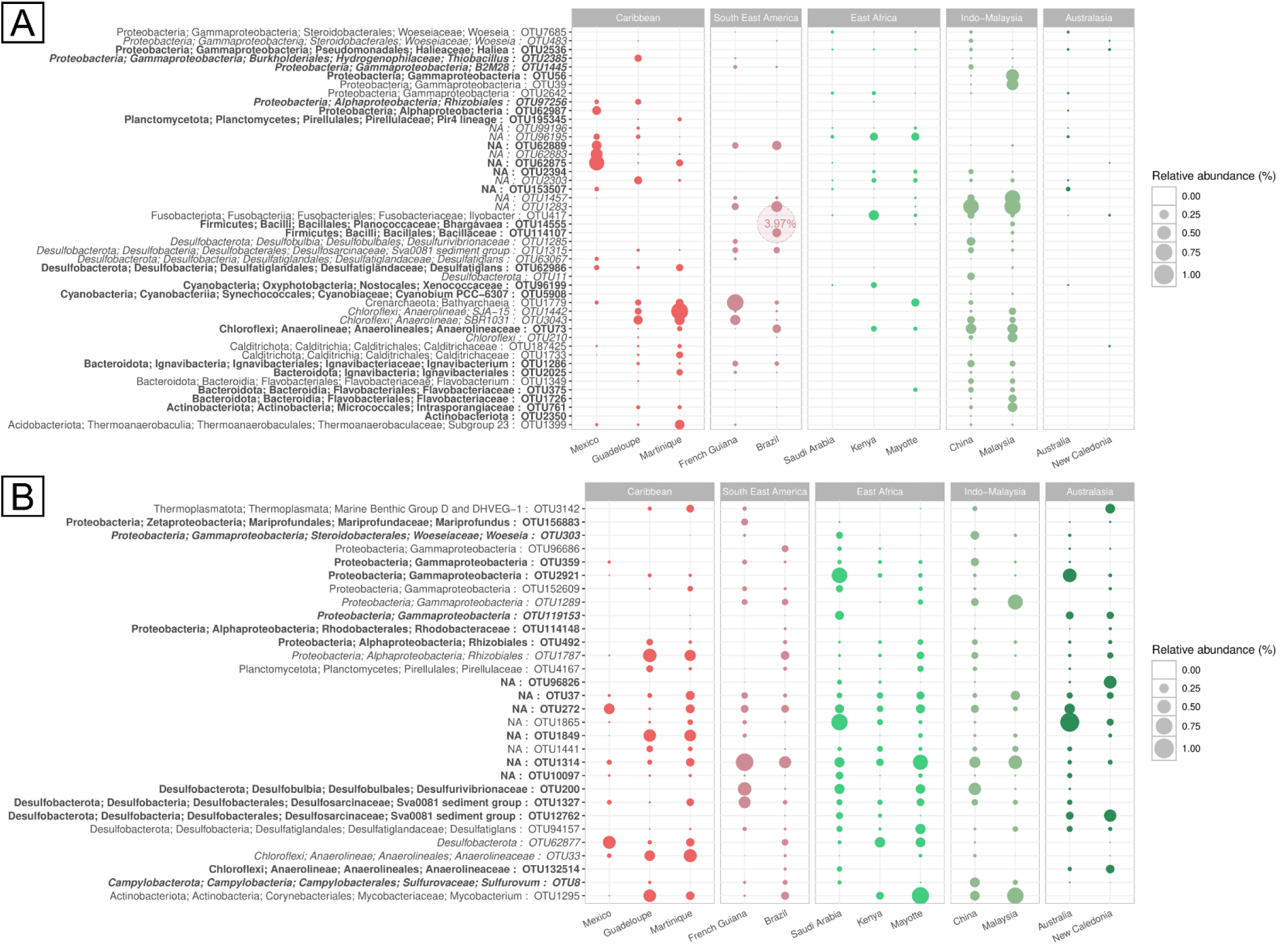
Relative abundances of taxa in each country, highlighting the 76 most important OTUs identified by the random forest model for the bioregion parameter. **(A) OTUs** not present in the core and **(B)** also present in the core. OTUs absent in the most important OTUs for the study ID model are shown in bold text, while OTUs present in the most important OTUs for the mangrove type model are italicized.

Overall, OTUs most representative of the Caribbean bioregion predominantly belonged to the classes *Anaerolineae* and *Calditrichaceae*. Southeast America was also represented by *Anaerolineae*; however, a notable distinction was observed in Brazil, where a *Bhargavaea* OTU stood out with a relative abundance of nearly 4%. This OTU, alongside members of the *Bacillaceae* family, were the only Firmicutes identified within the bioregion model results.

East Africa had fewer prominent OTUs compared to other bioregions, except for a *Xenococcaceae* OTU from the Cyanobacteria phylum. Nevertheless, this region displayed a fairly uniform representation of core OTUs. In contrast, almost all identified OTUs were present in the Indo-Malaysian bioregion, making it a highly representative area. Australasia had the fewest distinctive OTUs, suggesting that this bioregion was the least influential in classification decision-making.

No Archaeal OTUs were among the most important in the model, except for a Bathyarchaeia OTU. While this OTU was absent from the core, it remained significant within the study model and was predominantly found in the AEP.

## Discussion

### Microbial diversity hotspots and potential dispersal barriers in global mangrove sediments

Overall, a high averaged Shannon diversity index was found (6.7±0.5), which aligns with values reported by Lai *et al*. (2022) for mangrove sediment microbiota (5.7 to 11.1). This range is comparable to diversity levels found in the Earth Microbiome Project data for marine sediments, soil, and plant rhizosphere, and notably exceeds diversity found in other environments, such as water or animal host-associated microbiota (Bokulich *et al*., 2020).

Our findings indicate that microbial alpha-diversity is higher and more heterogeneous in the Indo-West Pacific (IWP) compared to the Atlantic-East Pacific (AEP), suggesting a potential microbial diversity hotspot in the Indo-Malaysian subregion, closely followed by East Africa. However, the observed nearly 1.5-fold difference in microbial alpha-diversity between AEP and IWP is less pronounced than the 3 to 6-fold disparity in mangrove tree diversity between these regions (Tomlinson, 2016). Previous studies have documented positive correlations between mangrove tree diversity and associated biota diversity (*e.g.* invertebrates and ichtyofauna) (Ellison et al. 1999; Reid et al., 2010; Lee et al. 2017), but our analysis found no significant relationship between microbial alpha-diversity and tree richness. On the whole, while the discrepancy in plant diversity likely arises from factors such as dispersal barriers, species-specific habitat preferences, and speciation rates, which have driven vicariance in mangrove flora (Duke et al., 2017), our results suggest a different dynamic for their associated microbial communities.

This absence of correlation between microbial and tree diversity contrasts with findings in other ecosystems, where microbial diversity often scales positively with plant diversity, especially in tropical regions (Liu et al., 2020). In terrestrial systems, this relationship may be driven by plant productivity, which shapes belowground niches and enhances microbial diversity (Zak et al., 2003). However, our results could point that mangrove prokaryotic communities are less subject to such selection compared to terrestrial tropical forests. Our findings suggest that mangrove microbiota may be less influenced by plant production dynamics than their terrestrial counterparts, indicating that other factors may govern microbial diversity patterns across mangrove bioregions. Future research should explore how local and regional environmental factors influence these global diversity patterns, particularly intercations between local tree assemblages and microbial communities.

### Latitudinal Diversity Gradients in Coastal and Estuarine Communities

We identified opposing latitudinal diversity gradients (LDGs) between coastal and estuarine mangroves. The negative microbial LDG in coastal mangroves mirrors the negative LDG seen in mangrove trees (Ellison et al., 2002) and associated ichthyofauna (Lee et al., 2017). This trend likely reflects increasingly unfavorable environmental conditions near 25°S and 25°N, including cooler temperatures and competition with salt marshes.

In contrast, in estuarine mangroves, microbial diversity increases with latitude, an unexpected finding that requires careful interpretation. The interaction between time and regional energy influences the rates of speciation, extinction or dispersal, however those mecanisms arehighly challenging to identify precisely (Cerezer et al., 2022; Saupe, 2023). Additionally, certain host-associated microbiomes have already been recognized as exceptions to the LDG theory without clear biotic or abiotic drivers currently explaining these patterns (Amaral-Zettler, 2021; Neu et al., 2021). Estuaries are subject to extent runoff and flooding, driven by the dynamics of climate and upstream watershed conditions. These factors alter the scales and rates of environmental variation, leading to differences in the drivers shaping microbial communities between estuaries and coasts (Adame et al., 2010).

This inverse microbial LDG in estuarine mangroves is another exception to the LDG theory, warranting further study on the biogeography and diversity of microbial communities in estuarian versus coastal environments.

### Distance-Decay Relationships in Mangrove Sediments

A significant, albeit weak, global microbial distance-decay relationship was detected. Distance-decay reflects how microbial compositional similarity declines with geographic distance, providing insights into dispersal barriers or the influence of environmental filtering across scales. Du *et al*. (2023) observed similar patterns in mangrove sediments within individual regions (e.g., China, South America). The strength of microbial distance-decay relationships can vary with spatial scale and environment type (Clark *et al*., 2021), with ecological divergences potentially masking biogeographical patterns (Härer and Rennison, 2022). Notably, parallels between microbial and macroorganisms may indicate shared environmental influences (Dickey *et al*., 2021). Therefore, the weakness of the global distance-decay relationship observed in the present study may reflect the confounding influence of global heterogeneities in mangrove ecosystems, that hampers its significance. Coastal mangroves communities, analyzed separately, exhibited a stronger distance-decay relationship (twice the slope and thrice the explanatory power). This suggests that shared ecological conditions, such as similar tidal influences or sediment charcateristics, drive consitent responses in both mangrove-asssociated microbes and their host vegetation. Conversely, the pooled global dataset likely encompasses a broader range of environmental conditions, diluting the distance-decay signal.

Multivariate analysis highlighted that estuarine mangrove communities were less dispersed than those of other mangrove types, possibly due to the more homogeneous conditions in estuaries. Such conditions include substantial freshwater inputs and reduced wave disturbances, which are known to enhance microbial diversity and impose stronger selective pressures on coastal sediment communities (Galand et al., 2016). Similarly in arid mangroves, Thomson et al (2022b) reported greater homogeneity in fringe mangroves due to water movements, while shrub mangroves displayed higher dispersion driven by the absence of such homogenizing factors.

Another plausible explanation for the weak global relationship observed is the influence of relic or dormant DNA. Relic DNA (extracellular or derived from non-intact cells) and dormant DNA (from inactive cells) are not subject to direct environmental selection (Carini *et al*., 2016; Locey *et al*., 2020). On average, approximately 33% of environmental prokaryotic DNA is relic, though this proportion can increase to 80% depending on the matrice characteristics (Lennon *et al*., 2018). While surface marine sediments are known to harbor significant quantities of relic DNA (Bairoliya et al., 2022), equivalent estimates for mangrove sediments remain limited.

Despite these complexities, the clear linear decline in community silmilarity with increasing spatial distance observed in coastal mangrove sediments underscores the presence of dispersal barriers for mangrove microbial communities. These findings highlight the nuanced interplay between environmental selection, dispersal, and biogeographical structure in mangrove ecosystems on a global scale.

### Global mangrove sediment core microbiota and bioregions representative taxa

At the global scale, microbial communities in mangrove sediments were predominantly composed of Gammaproteobacteria, which accounted for nearly one-sixth of all identified sequences. This finding highlights once again their ubiquity and ecological significance in mangrove ecosystems. As highly diverse bacterial class, Gammaproteobacteria are commonly found in marine environments, particularly in mangrove sediments and rhizosphere (Lai *et al*., 2022). They are involved in carbon, nitrogen and sulfur cycles (Dyksma et al., 2016; Yan et al., 2024), existing both as free-living forms and symbionts (Liang et al., 2007).

Among the Gammaproteobacteria, *Woeseia* was identified as the most abundant genus in the core microbiota, representing almost 11 % of the sequences. A recent study has suggested that *Woeseia* was a key player in nitrogen cycling within the surface sediments of Indo-Malaysian mangroves, where it expresses genes for ammonification, nitrification, and a truncated denitrification pathway (Qian et al., 2024). Additionally, several distinct OTU within *Woeseia* were identified as key bioregion and mangrove type markers through random forest (RF) models. These findings suggest that members of *Woeseia* exhibit biogeographical distribution of distinct ecotypes and niche differentiation, reflecting ecological specialization across bioregions.

The second most abundant phylum was Chloroflexi, primarily represented by family *Anaerolineaceae*, which was also part of the core microbiota. Within this family, *Litorilinea* accounted for 3.5% of total sequences. Known for their fermentative abilities, *Litorilinea* have been identified as keystone taxa in microbial interactions within mangrove sediments (Lin *et al*., 2018). Their fermentation products serve as substrates for sulfur-reducing bacteria (SRB), which preferentially utilize low-molecular-weight compounds as electron donors (Kristensen *et al*., 2008).

The third most abundant phylum, Desulfobacterota, includes SRB that are critical actors of sulfur cycling within mangrove ecosystems. With high oxygen demand and significant sulfate input from seawater, the sulfur cycle is estimated to account for approximately 50% of metabolic activity in mangrove sediments (Helfer and Hassenrück, 2021). Among the core microbiota, the genera *Desulfatiglans* and the Sva0081 sediment collectively representing nearly 6% of the total sequences. RF models identified specific OTU within this genera in the Caribbean, Southeast America, and Indo-Malaysia, aligning with findings from Du et al. (2023), who observed a higher prevalence of Desulfobacterota in South American mangrove sediments. This may be linked to higher organic carbon content, as SRB are known play key roles in organic matter mineralization. Spatial distribution of these OTU could therefore reflect variations in both the quantity and quality of organic carbon across bioregions. Additionnaly, variations in root morphology (e.g., *Avicennia* prop roots versus *Rhizophora* stilt roots) may influence SRB activity, with *Avicennia* promoting oxic zones that could inhibit SRB activity (Balk et al., 2016). Notably, the higher prevalence of *Avicennia* over *Rhizophora* in Australasia compared to the Caribbean for example, may help explain the absence of SRB in Australasia in the RF analysis. Concerning those samples, Luis et al. (2019) observed significantly higher redox potentials under *A. marina* (-136.9 to 74.4 mV) compared to *R. stylosa* (-176.6 to -72.8 mV) in New Caledonia, while Fiard et al. (2024) reported more reduced conditions in Martinique, ranging from -200 mV and -318.3 ± 55.7 mV.

The Desulfobacterota phylum also includes sulfide-oxidizing bacteria (SOB), which are frequently found in anoxic layers with thin aerobic zones, such as the top few millimeters of sediments. These bacteria contribute maintaining sediment homeostasis (SamKamaleson and Gonsalves, 2019). Among SOB, cable bacteria from the *Desulfobulbaceae* family, such as *Desulfobulbus*, were part of the core microbiota. Unique in their ability to transport electrons over centimetre-scale distances, cable bacteria link oxic layers to deeper organic-rich sediments (Pfeffer *et al*., 2012; Burdorf *et al*., 2017). Their distribution is strongly influenced by bioturbation, as sediment-disturbing fauna can serve these conductive cables (Malkin *et al*., 2014). Bioturbation rates, which vary between 20 to 80% of mangrove sediments depending on the mangrove type and region (Booth *et al*., 2019), could play a critical role in determining cable bacteria prevalence. However, SOB did not appear in the RF analysis, precluding any clear identification of biogeographical patterns. SOB are also represented within Gammaproteobacteria, with the phototrophic genus *Thiogranum* identified as part of the core microbiota, represented almost 4% of the sequences.

The archaeal component of the core microbiota was dominated by Crenarchaeota, although one *Bathyarchaeia* OTU appeared in the RF results, mainly in the AEP regions. *Bathyarchaeia* are anaerobes capable of degrading complex organic molecules like lignin, contributing to the breakdown of plant-derived compounds (Yu et al., 2018). Methanogens, represented mainly by *Methanomassiliicoccales* from the *Thermoplasmatota* phylum, were scarce, likely due to the shallow sediment depth (20 cm) analyzed in this study. Methanogens typically thrive in deeper, strictly anoxic sediments where competition with SRB is reduced (Kristensen et al., 2008).

A surprising observation was the disproportionately high abundance of Firmicutes in Brazilian mangroves. An OTU from the Firmicutes phylum, assigned to the *Planococcaceae* family, was identified in the RF analysis, indicating its key biogeographic importance across bioregions. Firmicutes dominance was previously noted in sediments of South American mangroves by Lai *et al*. (2022). Fernández-Cadena *et al*. (2020) even reported *Planococcaceae* in their top 50 most abundant OTUs in Ecuadorian mangroves. However, this OTU was also prevalent in Malaysian mangroves (Mai *et al*., 2021), and comparable patterns have been observed in India (Kutty *et al*., 2023), suggesting that their distribution cannot be solely explained by dispersal limitations. Instead, environmental factors likely play a significant role. As k-strategists, Firmicutes are well-adapted to water stress through mechanisms such as thick cell walls and endospores formation, as seen in *Planococcaceae* representatives (Pan et al., 2022; Shivaji et al., 2014). Further research is needed to understand their responses to spatial and temporal stressors in different geomorphological mangroves, particularly across wet and dry periods.

In summary, the biogeographical patterns of mangrove sediment microbiota reflect intricate ecological interactions influenced by environmental heterogeneities, root morphology, and metabolic processes. Further investigations, particularly those integrating metadata such as carbon content and root proximity, are essential to disentangle the complex dynamics shaping microbial distributions across mangrove bioregions.

### Challenges and perspectives for global-scale analyses in mangrove sediments

Our study revealed that approximately 23% of OTUs remained unassigned at the phylum level, consistent with findings from other studies not included in this meta-analysis (Haldar and Nazareth, 2018; Kutty *et al*., 2023; Yun *et al*., 2017). This significant proportion of “microbial dark matter” highlights the vast reservoir of uncharacterized microbial diversity in mangrove sediments, a phenomenon common in marine sediments and extreme environments (Schultz *et al*., 2023). Notably, some of these unassigned sequences were among the most ubiquitous in our dataset, including in the core microbiota, suggesting that they may hold critical insights into mangrove microbial biogeography, as suggested by their significance in the bioregion RF model. Advances in long-read sequencing technologies (Tedersoo et al 2021) may enhance taxonomic resolution, however database limitations persist due to the difficulty of isolating and identifying microorganisms from the complex, organic-rich environment of mangrove sediments.

Another key observation was the notable inter-study “patchiness”, characterized by limited overlap in OTU across studies. This variability was evident in the beta-diversity analyses, where the “study” variable accounted for a significant portion of the observed variation. For instance, East Africa and Australasia exhibited the highest overlap among all bioregions, sharing 6,419 OTUs exclusively, with closely intertwined NMDS plots . This overlap was largely attributed to the fact that these samples, derived from arid mangroves, were collected and analyzed within the same study (Thomson *et al*., 2022a). This “patchiness” likely reflects the high sensitivity of microbial communities to the pronounced heterogeneity of mangrove habitats, shaped by diverse microhabitats like fauna burrows, roots, litter, water channels, and pools (Nagelkerken *et al*., 2008), as well as ecological and geomorphological types (fringe, riverine, inland, shrub, basin, lagoon, estuary, delta, oceanic island, and coastal mangroves; Twilley *et al*., 2019). Additional drivers of microbial variability include climatic differences, anthropization impacts, mangrove age, or mangrove species composition, all of which influence microbial diversity and activity (Gómez-Acata, Teutli *et al*., 2023; Fiard *et al*., 2022 ; Muwawa *et al*., 2021). This inherent complexity makes universal sampling strategies in mangrove studies particularly challenging.

Despite efforts to standardize sampling strategies (*e.g.,* sediment depth, native mangrove ecosystems), unavoidable field-level variability and methodological inconsistencies -such as differences in sample preservation and storage, DNA extraction methods, sequencing platforms, primers, targeted region, and sequencing runs-conbtributed to inter-study variability (Compson *et al*., 2020; Salter *et al*., 2014; Abellan-Schneyder *et al*., 2021). As noted by Du *et al*. (2023), using different bioinformatics pipelines for 16S rRNA metabarcoding presents some challenges for comparing mangrove data on a global scale, but it also opens up opportunities for more in-depth future analyses. Consequentely, while disentangling ecological factors from methodological biases remains a challenge in microbial ecology, it also presents an opportunity to improve the accuracy of microbial ecology research .

To address these challenges and drawing inspiration from other successful initiatives like the *Earth Microbiome Project* (Shaffer *et al*., 2022), the *Mangrove Microbiome Initiative* (Allard *et al*., 2020) could propose harmonized sampling and sequencing methodologies, facilitating more consistent comparisons of mangrove microbial communities worldwide. Such efforts would enhance our ability to resolve global patterns and underlying drivers of microbial communities in mangrove ecosystems.

## Conclusion

This meta-analysis provides novel insights into the global biogeography of mangrove sediment microbiota, uncovering patterns of diversity, dispersal, and community composition that highlight the ecological complexity of these ecosystems. The identification of microbial diversity hotspots in the Indo-West Pacific and the clear differences between coastal and estuarine communities underscore the intricate biogeographical dynamics at play. While our findings confirm the presence of dispersal barriers and distance-decay relationships in coastal mangroves, they also suggest that environmental factors, rather than plant diversity alone, are key determinants for shaping prokaryotic community structures. The challenges posed by inter-study variability, methodological discrepancies, and “microbial dark matter” amphasize the need for standardized methodsin future research. Despite these limitations, our findings lay a foundational framework for understanding global microbial biogeography in mangroves, paving the way for further inititiaves. Moving forward, coordinated efforts like the *Mangrove Microbiome Initiative* (Allard et al., 2020) could unlock a deeper understanding of the functional roles and ecological processes governing mangrove microbiomes.

## Data availability statement

Data are available in the SRA of NCBI following the accession numbers listed in the bioproject column of Table 1. All scripts have been deposited in Zenodo for public access (https://doi.org/10.5281/zenodo.13991716).

## Acknowledgements

We thank everyone who responded to our request regarding missing data and kindly provided additional information, including, in no particular order, Luisa Falcón, Meng Li, Zhichao Zhou, Prinpida Sonthiphand, Timothy Thomson, Weidong Chen, Donghui Wen, Jenny Booth, and Benjamin Wainwright. Special thanks to Timothy Thomson, who also graciously agreed to meet virtually for enriching scientific discussions. We also acknowledge the valuable advice of David Nerini on constructing random forest models. This work is part of the Multidisciplinary Thematic Network “RTP Mangroves” (CNRS), which unites French research expertise on mangroves from various laboratories.

