## Supplementary figures for "Global Biogeography of Prokaryotes in Mangrove Sediments: Spatial Patterns and Ecological Insights from 16S rDNA Metabarcoding"

### SUPPLEMENTARY MATERIALS

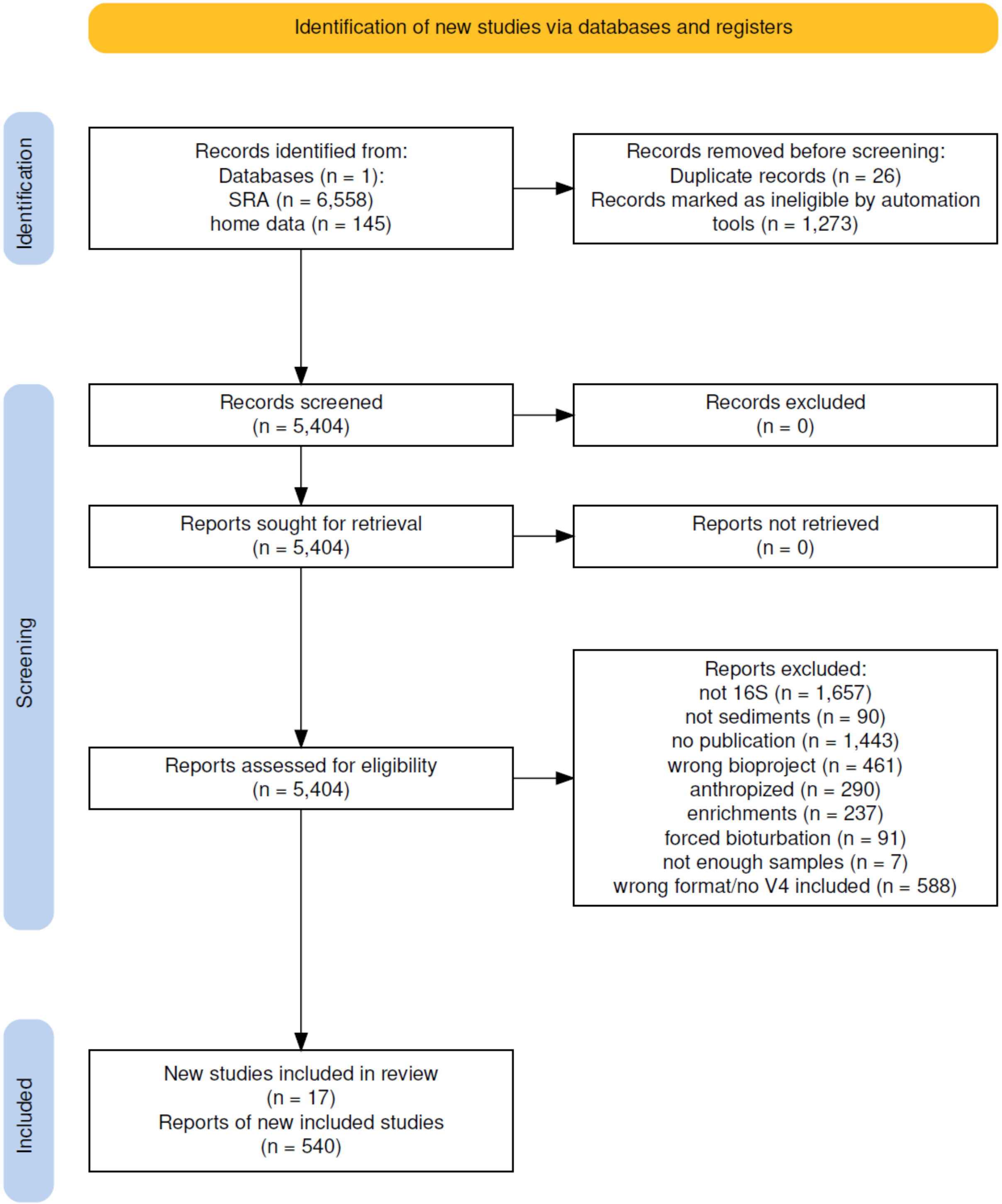


**Supplementary Figure 1.** PRISMA diagram outlining of the data collection process for the meta-analysis. Reports refer to fastq files of 16S metabarcoding of individual environmental samples. Studies correspond to Bioprojects under which these files were deposited.

**Supplementary Table 1.** Summary of Bioprojects IDs, primer pairs, and associated metadata used in the meta-analysis.

*see excel file*

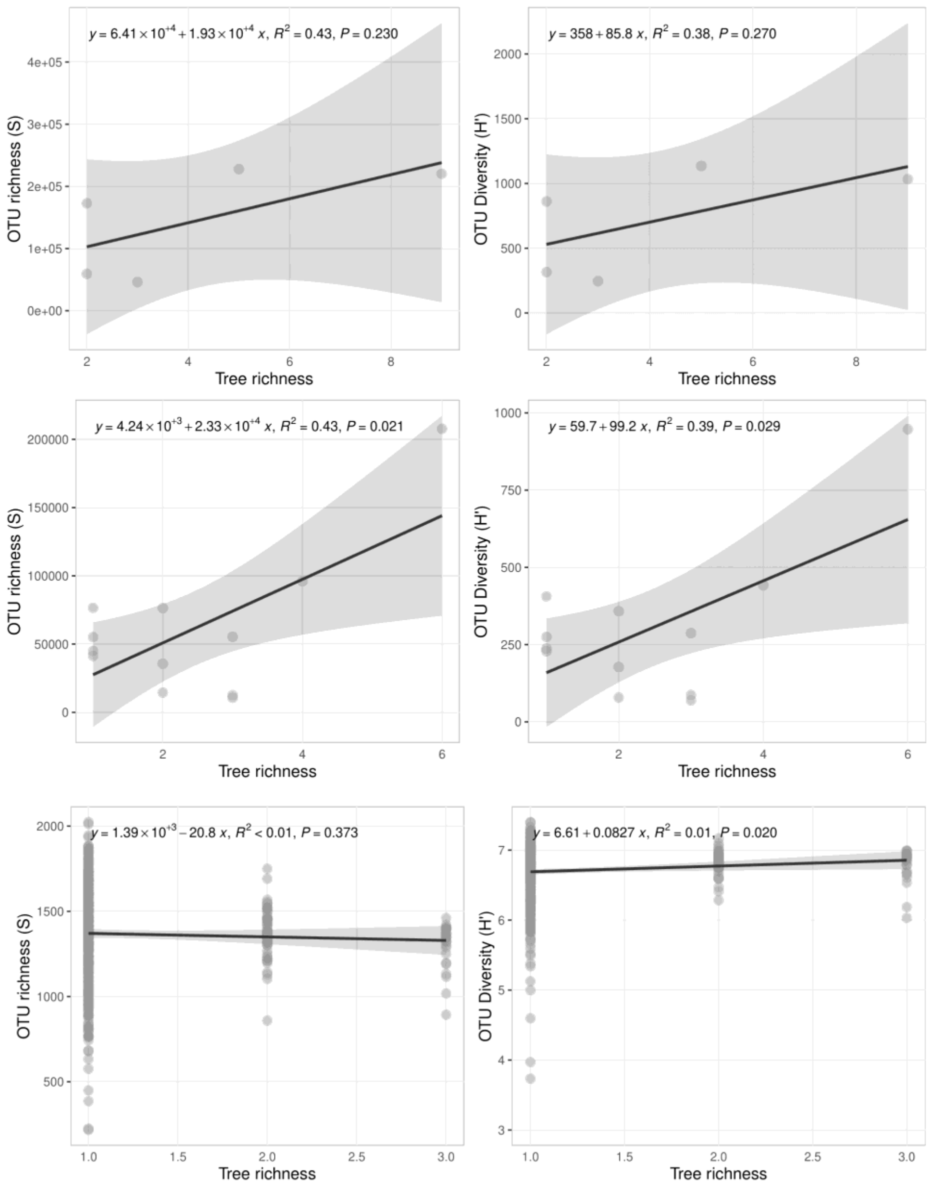


**Supplementary Figure 2.** Linear regressions showing the absence of relationship between OTU alpha-diversity and tree species richness. Analyses were conducted at three scales: bioregion (top), country (middle) and sample level without pooling (bottom).

### *Functional diversity*

Putative functional group profiles were assigned to the rarefied OTU table (excluding singletons), based on the SILVA 138.1 taxonomy. The analysis utilized the 'collapse_table.py' script by Louca *et al.* (2016), which integrated the Functional Annotation of Prokaryotic Taxa (FAPROTAX) database v1.2.10. The FAPROTAX database is a curated resource comprising peer-reviewed references on known microbial metabolisms and their associated clades.

Louca, S., Parfrey, L. W., & Doebeli, M. (2016). Decoupling function and taxonomy in the global ocean microbiome. *Science*, *353*(6305), 1272-1277.

Functional groups could be assigned to 13.6% of total OTUs, representing 68 out of the 92 groups in the database. High heterotrophy characterized the mangrove sediments, including both aerobic and anaerobic pathways (Supplementary Figure 4). Aerobic respiration was the most prevalent function, comprising 23% of the assigned OTUs. These were linked to 28 genera, several of which overlapped with the OTU-based core, including *Lewinella*, *Pseudomonas*, *Robiginitalea*, *Vibrio*, *Haliscomenobacter*, *Blastopirellula*, *Rhodopirellula,* as well as members of the families *Cyclobacteriaceae* and *Sphingomonadaceae*.

Sulfur metabolism played a significant role, with sulfur-related functional groups expressed ubiquitously across all bioregions. Sulfur compounds were used either as electron acceptors (respiration) or electron donors (oxidation). Notably, 28% OTUs were involved in sulfur compounds respiration, largely attributed to *Desulfosarcinaceae*, *Desulfatiglandaceae* and *Desulfobacteraceae*. Core genera, including *Sva0081 sediment group*, *Desulfobulbus*, *Desulfobacca*, *Desulfatiglans,* and *Desulfopila*, underscored the importance of sulfate respiration in mangrove sediments. Dark sulfur oxidation and anoxygenic phototrophic sulfide oxidation accounted for 1.3% and 1.6% of OTUs respectively. Key taxa included *Sulfurimonas* (Campylobacterota) and *Magnetospira* (Alphaproteobacteria) for dark oxidation, as well as *Halochromatium*, *Thiogranum* and other *Ectothiorhodospiraceae* (Gammaproteobacteria) for anoxygenic photoautotrophy. Oxygenic photoautotrophy while represented by only 4.3% of OTUs, was one of the most abundant functions, being exclusively attributed to Cyanobacteria.

The genus *Cyanobium PCC-6307* (Synechococcales) was the sole representative of the OTU-based core. Interestingly, OTUs related to human-associated taxa were absent from the OTU-based core, suggesting minimal anthropogenic microbiota influence on mangrove microbial communities. Other interesting functional groups related to nitrogen cycle, methanogenesis, plant composites decomposition (cellulolysis, xylanolysis) or endofauna exosqueletons decomposition (chitinolysis) were also assigned. Overall, the Indo Malaysia bioregion exhibited the highest functional richness, as confirmed by the Dunn's Test ($p < 0.001$ for each paired comparison).

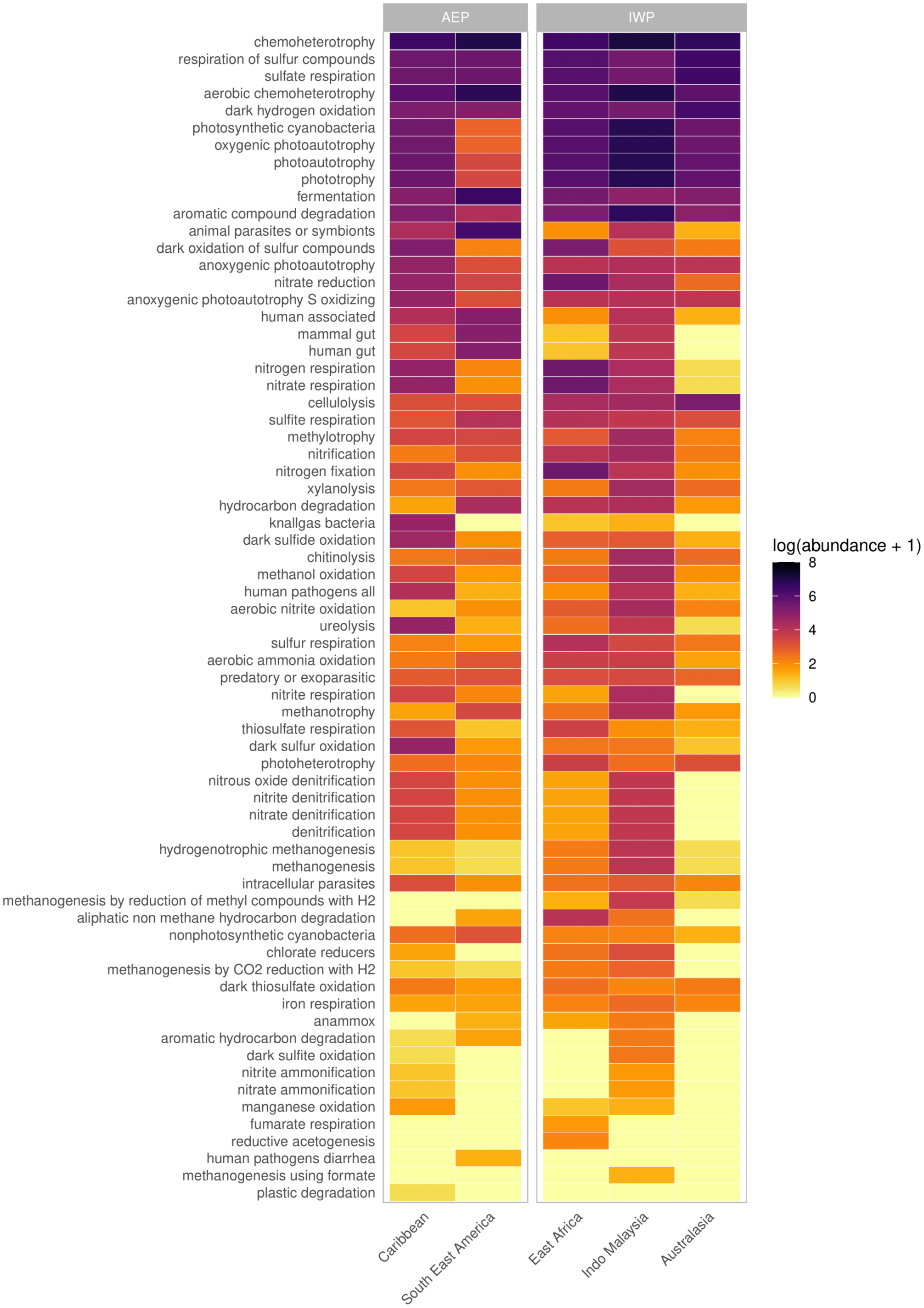


**Supplementary Figure 3.** Heatmap illustrating ranked relative abundances of taxonomically assigned functions across bioregions, based on the FAPROTAX database. Data were derived from the pooled samples included in the meta-analysis.

**Supplementary Table 2.** Results of PERMANOVA analyses assessing the Jaccard distance of OTUs across various class parameters.

| | df* | SS* | $R^2$ | Pseudo-F | P-value | |
|---|---|---|---|---|---|---|
| **Bioregion** | 4 | 20,701 | **0,084** | 12,249 | 0,001 | *** |
| **Residual** | 535 | 226,043 | 0,916 | | | |
| **Total** | 539 | 246,744 | 1,000 | | | |
| **Mangrove type** | 3 | 16,844 | **0,068** | 13,090 | 0,001 | *** |
| **Residual** | 536 | 229,900 | 0,932 | | | |
| **Total** | 539 | 246,744 | 1,000 | | | |
| **Koppengeiger classification** | 4 | 24,374 | **0,099** | 14,660 | 0,001 | *** |
| **Residual** | 535 | 222,370 | 0,901 | | | |
| **Total** | 539 | 246,744 | 1,000 | | | |
| **Sampling year** | 5 | 24,110 | **0,098** | 11,566 | 0,001 | *** |
| **Residual** | 534 | 222,633 | 0,902 | | | |
| **Total** | 539 | 246,744 | 1,000 | | | |
| **Extraction kit** | 5 | 19,988 | **0,081** | 9,414 | 0,001 | *** |
| **Residual** | 534 | 226,755 | 0,919 | | | |
| **Total** | 539 | 246,744 | 1,000 | | | |
| **Sequencing platform** | 1 | 7,463 | **0,030** | 16,780 | 0,001 | *** |
| **Residual** | 538 | 239,281 | 0,970 | | | |
| **Total** | 539 | 246,744 | 1,000 | | | |
| **Forward primer** | 4 | 22,934 | **0,093** | 13,705 | 0,001 | *** |
| **Residual** | 535 | 223,810 | 0,907 | | | |
| **Total** | 539 | 246,744 | 1,000 | | | |
| **Reverse primer** | 4 | 20,930 | **0,085** | 12,397 | 0,001 | *** |
| **Residual** | 535 | 225,814 | 0,915 | | | |
| **Total** | 539 | 246,744 | 1,000 | | | |
| **Taq polymerase** | 8 | 28,284 | **0,161** | 9,087 | 0,001 | *** |
| **Residual** | 379 | 147,460 | 0,839 | | | |
| **Total** | 387 | 175,744 | 1,000 | | | |
| **Study ID** | 16 | 64,913 | **0,263** | 11,669 | 0,001 | *** |
| **Residual** | 523 | 181,831 | 0,737 | | | |
| **Total** | 539 | 246,744 | 1,000 | | | |

**Supplementary Table 3.** Proportions of unassigned sequences between studies and across taxonomic levels, illustrating variations in assignment consistency within the dataset.

| Country | Study | Kingdom | Phylum | Class | Order | Family | Genus | Species |
|---|---|---|---|---|---|---|---|---|
| Mexico | PRJNA550111 (Gómez-Acata, Teutli et al 2023 | 1,88% | 31,68% | 40,75% | 57,91% | 76,11% | 89,72% | 99,94% |
| Guadeloupe | PRJNA1118708 | 1,02% | 27,11% | 32,43% | 48,85% | 62,28% | 83,46% | 99,83% |
| Martinique | PRJNA874590 (Fiard et al 2024) | 0,70% | 27,92% | 32,15% | 49,34% | 63,19% | 84,05% | 99,91% |
| French Guiana | PRJNA735070 (Fiard et al 2022) | 0,73% | 24,52% | 30,22% | 51,50% | 65,37% | 84,99% | 99,92% |
| Brazil | PRJNA608697 (de Santana et al 2021) | 0,04% | 18,55% | 25,26% | 44,12% | 57,55% | 79,28% | 99,90% |
| Brazil | PRJNA817610 (Machado da Costa et al 2023) | 0,20% | 20,78% | 26,29% | 42,72% | 57,13% | 77,81% | 99,26% |
| Mayotte | PRJNA1118920 | 0,10% | 27,53% | 30,59% | 51,23% | 63,96% | 84,92% | 99,94% |
| Kenya | PRJNA644929 (Muwawa et al 2021) | 0,00% | 35,04% | 38,64% | 57,57% | 68,60% | 83,27% | 99,87% |
| Saudi Arabia | PRJNA720541 (Thomson et al 2022) | 0,89% | 25,57% | 31,80% | 50,55% | 64,59% | 83,58% | 99,97% |
| China | PRJNA732523 (Liu et al 2023) | 0,08% | 20,51% | 29,33% | 48,35% | 63,10% | 81,59% | 99,94% |
| China | PRJNA475455 (Zhang et al 2019) | 1,20% | 24,96% | 32,74% | 51,48% | 67,80% | 85,18% | 99,89% |
| China | PRJNA797991 (Zhang et al 2022) | 0,65% | 24,20% | 31,26% | 52,99% | 67,59% | 85,69% | 99,82% |
| China | PRJNA404001 (Zhu et al 2018) | 0,00% | 18,19% | 21,98% | 37,68% | 46,98% | 72,27% | 99,58% |
| China | PRJNA779243 (Zhu et al 2022) | 0,01% | 20,48% | 27,07% | 44,97% | 58,97% | 79,99% | 99,60% |
| Malaysia | PRJNA756333 (Mai et al 2021) | 0,00% | 17,47% | 22,85% | 40,43% | 55,10% | 76,77% | 99,40% |
| New Caledonia | PRJEB25548 (Luis et al 2019) | 0,19% | 22,76% | 27,14% | 49,11% | 66,84% | 87,37% | 100,00% |
| Australia | PRJNA720541 (Thomson et al 2022) | 0,89% | 25,57% | 31,80% | 50,55% | 64,59% | 83,58% | 99,97% |

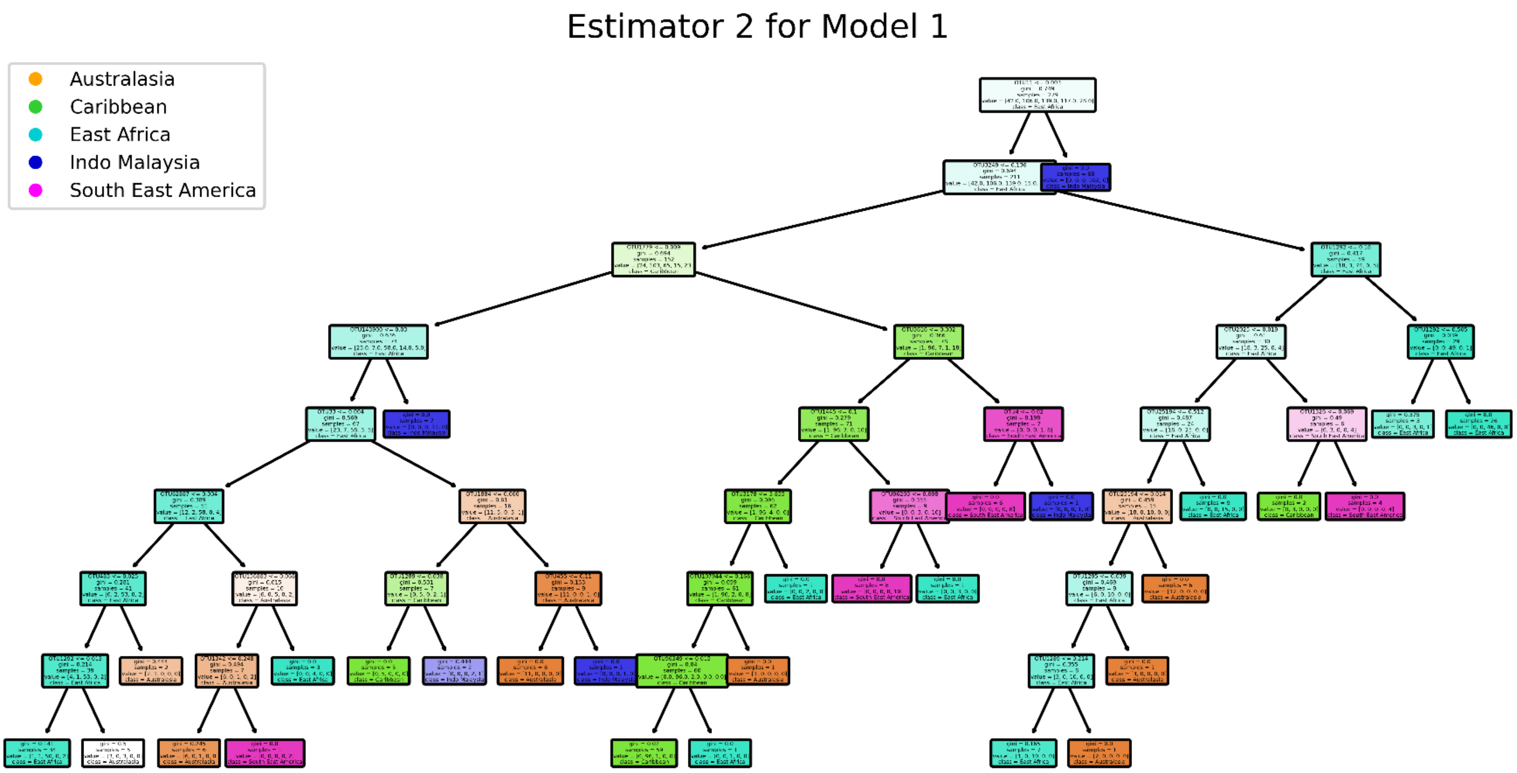


**Supplementary Figures 4.** Example of a random forest model trees (estimators) generated for the bioregion class. Only one tree is shown here for visualization; additional models can be found in the supplementary files.

**Supplementary Table 5.** Results of random forest models, including scores and the most important OTUs associated with each class.

*See excel file*
